# Rapid *in vitro* synthesis of DNA templates via Sidewinder for polyadenylated Hantavirus mRNA vaccine candidates

**DOI:** 10.64898/2026.05.22.727328

**Authors:** Ezri Abraham, Joel Andrade, Alexander Davis, Tomasz Gawda, Jacob Glanville, Daniel Graves, Jianyi Huang, Jeongsoo Hur, Sangil Kim, Jean-Sebastien Paul, Noah Evan Robinson, Charles Sanfiorenzo, Sixiang Wang, Raymond J Zhang, Weilin Zhang, Tianhua Zhao, Jie Zhou, Kaihang Wang

**Author notes:** Department of Chemical Engineering, Stanford University, Stanford, CA 94305 USA. These authors contributed equally to this work, ordered alphabetically by last name (Ezri Abraham, Joel Andrade, Alexander Davis, Tomasz Gawda, Jacob Glanville, Daniel Graves, Jianyi Huang, Jeongsoo Hur, Sangil Kim, Jean-Sebastien Paul, Noah Evan Robinson, Charles Sanfiorenzo, Sixiang Wang, Raymond J Zhang, Weilin Zhang, Tianhua Zhao, Jie Zhou).

## Abstract

As the recent COVID-19 pandemic illustrated, zoonotic viruses and other pathogens pose a credible threat to public health. Recent advancements in vaccine technology, particularly mRNA vaccines, provide key tools for an effective and swift public health response. Although mRNA vaccines can be developed more quickly than traditional vaccines, fast and accurate construction of DNA templates for these vaccines remains a critical bottleneck. Using our novel DNA assembly technology, Sidewinder, we rapidly designed and built multiple mRNA vaccine candidates to guard against a potential outbreak of Hantavirus (ANDV). We successfully constructed the DNA templates from oligo pools and produced the mRNA for three vaccine candidates in just 2 days after delivery of the synthetic DNA oligos.

## Introduction

Rapid and robust vaccine production has attracted significant attention, particularly since the outbreak of SARS-CoV-2 in the COVID-19 pandemic. Among the various vaccine platforms, messenger RNA (mRNA) vaccines have demonstrated remarkable efficacy, safety, and adaptability against infectious pathogens^1^. Producing the RNA payload in the mRNA vaccine relies on the production of DNA which serves as the template for *in vitro* transcription (IVT)^2^. The rapid construction of viral antigenic DNA sequences, particularly those which are long and highly complex, remains technically challenging as conventional DNA assembly methods often depend on multiple *in vivo* cloning steps, assembly-constrained sequence optimization, and labor-intensive procedures that can compromise speed, efficiency and reliability^3,4^. Such limitations become acutely apparent during outbreaks of transmissible pathogens, where slow time-to-construction of DNA templates for mRNA vaccines can substantially delay public health response.

On May 2nd, 2026, the World Health Organization (WHO) announced that Andes virus (ANDV), a strain of Hantavirus, caused 3 deaths on a cruise ship that left from Argentina^5^ (**Fig. 1a**). As of May 13th, the virus had infected 11 total people, all of which were on the cruise ship^6^. While this virus is not reported to be spreading to those who were not on the cruise, a study of a previous ANDV outbreak reported a super-spreading capability of the virus if no countermeasures are applied^7^.

**Fig 1.**
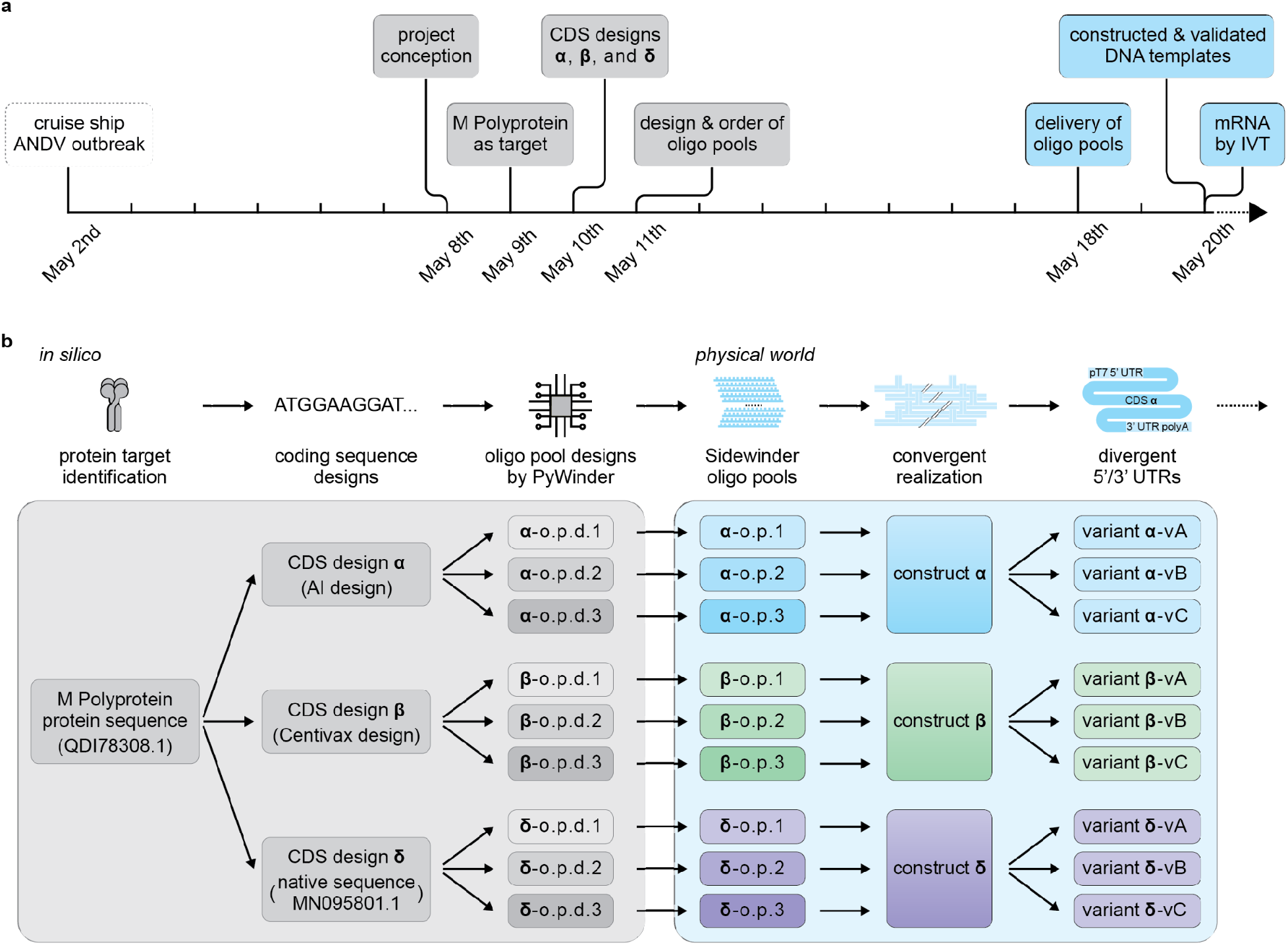
Sidewinder enables failsafe parallel design and construction of DNA templates for mRNA vaccine candidates. **a)** Timeline depicts the sequence of events between the ANDV outbreak, the conceptualization of the project, through to the expression of mRNA. Events depicted in gray are *in silico* and events depicted in color are experimental with just 2 days between receiving the synthetic DNA oligos and construction and *in vitro* transcription (IVT) of the 3.75 kb mRNA construct. **b)** Schematic flowchart overview depicting the multifaceted approach taken to design, DNA template construction leading to multiple outputs for downstream characterization. From a single protein target, CDS α, β, and δ are designed as different coding sequences to encode the same amino acid sequence (Genbank accession: QDI78308.1). For each CDS, three oligo pool designs (o.p.d.) are generated using PyWinder which when assembled, each construct the exact sequence of the parent CDS. All 9 o.p.d. are ordered and delivered as oligo pools (o.p.) which are all processed simultaneously to increase likelihood of success. Each and all of the 9 o.p. reconstitute the corresponding construct α, β, or δ as a physical molecule that is then processed into multiple downstream pipelines for different applications (variants vA, vB, vC).

On May 8th, we conceptualized a project to produce the DNA templates for ANDV mRNA vaccine candidates to prepare for a potential epidemic of ANDV^8^ (**Fig. 1a**). We recently developed a novel approach to DNA assembly, Sidewinder, which is capable of rapidly constructing large and complex DNA molecules from individual oligos^9^ (6) or oligo pools^10^ (7). Sidewinder has been demonstrated to efficiently assemble large constructs up to 12.5 kb entirely *in vitro* using oligo pools and has measured misconnection rates as low as 1 in 10,000,000, making it the most accurate DNA assembly method available^10^ (7). The speed, reliability, and accuracy of Sidewinder positions it as the ideal method for DNA construction, and in particular for the rapid construction of DNA templates for mRNA vaccines entirely *in vitro*.

In this current work we aimed to demonstrate how rapidly we could develop DNA templates for mRNA vaccine candidates for ANDV using Sidewinder. We first designed multiple vaccine candidate sequences expressing ANDV envelope glycoproteins and used our novel oligo design software PyWinder^10^ (7) to design oligos for Sidewinder assembly. After the oligo pools arrived, we constructed the 3.75 kb DNA template entirely *in vitro*. Finally, we used PCR to append the template for the 3’ poly(A) tail and performed *in vitro* transcription to create mRNA for ANDV vaccine candidates.

On May 18th, we received the delivery of synthetic DNA oligo pools (**Fig. 1a**). On May 20th,we finished construction and validation of DNA templates and *in vitro* transcribed them into mRNA for downstream vaccine development.

## Results

### Failsafe parallel design of mRNA vaccine candidates

mRNA vaccines require the construction of a synthetic DNA molecule as a template prior to mRNA production. To ensure a short time-to-production of mRNA and to eliminate any mission-critical failures, we took a multifaceted approach to construct the physical DNA molecules for multiple Hantavirus ANDV mRNA vaccine candidates. Towards this end, this study presents a rapid and robust workflow for DNA template construction for mRNA production with minimal time between design, construction, and *in vitro* transcription of capped, polyadenylated mRNA.

Drawing inspiration from the rapid response and production of mRNA vaccines for the COVID-19 pandemic^1^, we conceptualized our project for constructing mRNA with intended use in an mRNA vaccine against ANDV on May 8th (**Fig. 1a**). We ordered the short synthetic single stranded DNA “oligos” for DNA construction on May 11th which arrived on May 18th (**Fig. 1a**). Owing to our novel methodology for DNA assembly, Sidewinder^6^, we successfully *in vitro* transcribed mRNA on May 20th within just 2 days of acquiring the synthetic oligos (**Fig. 1a**).

To achieve this, at a high level, we *in silico* identified the antigen target, designed multiple coding sequences (CDS), and designed multiple Sidewinder reactions for each CDS (**Fig. 1b**). Then *in physico* we received the physical synthetic DNA oligos, processed each unitary oligo pool with Sidewinder assembly to converge each pool back to a CDS design as a physical DNA molecule, and processed the construct into multiple variants for differing downstream applications (**Fig. 1b**). To increase our likelihood of success, we designed a multifaceted approach with intentionally built-in failsafe redundancies at multiple stages (**Fig. 1b**). Owing to the ease and efficiency of the Sidewinder assembly method, all avenues were pursued simultaneously with minimal increased labor due to parallel processing of unitary oligo pools.

The overall design of our construct for mRNA production consists of five major components: a T7 promoter, a 5′ untranslated region (5′ UTR), an antigen CDS, a 3′ untranslated region (3′ UTR), and a poly(A) tail. We first identified the protein target for the mRNA vaccine which will compose the CDS component. ANDV encodes 4 proteins: a glycoprotein precursor (hereafter referred to as “M Polyprotein”), the nucleocapsid protein, an RNA-dependent RNA polymerase, and a putative non-structural protein^11^. The M Polyprotein (Genbank Accession: QDI78308.1) undergoes cleavage into two surface glycoproteins, Gn and Gc, via the host signal peptidase complex in the endoplasmic reticulum, and is responsible for viral entry into host cells^12^. As surface glycoproteins have been recognized as good mRNA vaccine targets^13^, our industrial collaborator Centivax identified the M Polyprotein for our mRNA vaccine candidates. We further analyzed its potential for MHC Class I and II presentation to CD8^+^ Cytotoxic T-cells by nucleated cells and CD4^+^ Helper T-cells by antigen-presenting cells respectively^14,15^, and found M polyprotein can be cleaved into peptides that have a high likelihood for both (**Methods**).

We then generated *in silico* three CDS designs (alpha (α), beta (β), and delta (δ)) with alternative coding choices from this amino acid sequence (**Fig. 1b**). The three CDS designs function as mutual failsafes in the event any failure points arise in oligo synthesis, DNA construction, DNA amplification, RNA transcription, or RNA secondary structures. Further, codon choice is known to affect transcription and translation^16,17^ potentially influencing large scale production or immunogenicity in the downstream development of the vaccine^18-20^.

The CDS design α was generated using the generative mRNA vaccine design and optimization framework GEMORNA, which has previously been shown to enhance mRNA stability and translational efficiency^21^. The CDS design β was optimized independently by our industrial collaborator, Centivax. The CDS design δ was the native viral coding sequence without optimization (Genbank accession: MN095801.1). The 5′ UTR and 3′ UTR of all three vaccine candidates were designed using GEMORNA.

After completing the CDS designs for α, β, and δ, we designed the synthetic oligos for the construction of the physical 3.75 kb synthetic DNA construct. The oligos were designed for Sidewinder DNA assembly: a technique which uses barcodes and a unique three-way junction (3WJ) structure to achieve high-fidelity DNA assembly with a measured misconnection rate of 1 in 10,000,000 at the assembly junction^7^. This high specificity of Sidewinder as well as the ease of processing multiple constructs simultaneously from unitary oligo pools enabled CDS designs α, β, and δ to each be constructed three independent times using differing Sidewinder oligo pool designs (o.p.d.) for a total of 9 candidate pools. Our novel design algorithm PyWinder^10^ was used to generate the three o.p.d. for each sequence with expanded thermodynamic filtering and expanded Sequence Search Minimization barcode set generation (**Methods**) (**Fig. 1b**).

### Failsafe parallel construction of mRNA vaccine candidates

Upon the ordering of the o.p.d. (α_o.p.d 1_, α_o.p.d 2_, α_o.p.d 3_, β_o.p.d. 1_, β_o.p.d. 2_, β_o.p.d. 3_, δ_o.p.d.1_, δ_o.p.d.2_, and δ_o.p.d.3_) on May 11th, the physical oligo pools (o.p.) arrived midday May 18th (**Fig. 1a)**. Each o.p. contained 138 oligos, encoding for a 69 fragment assembly composing the corresponding 3.75 kb target CDS. Within each o.p., all fragments were annealed and assembled in parallel in the same reaction tube composing the 3WJ assembly intermediate (**Fig. 2a**). Using the 3WJ assemblies as template, five intermediate amplicons were generated from each pool that were approximately 0.7-1 kb in length and together span the length of the ~4 kb target construct. Notably, all 45 intermediate amplicons produced a prominent band of the correct size (**Fig. 2b**). There are slight variations in band intensity or, in some cases, the presence of minor PCR byproducts such as primer dimers or mispriming for the intermediate amplicons across CDS designs and o.p.d.

**Fig 2.**
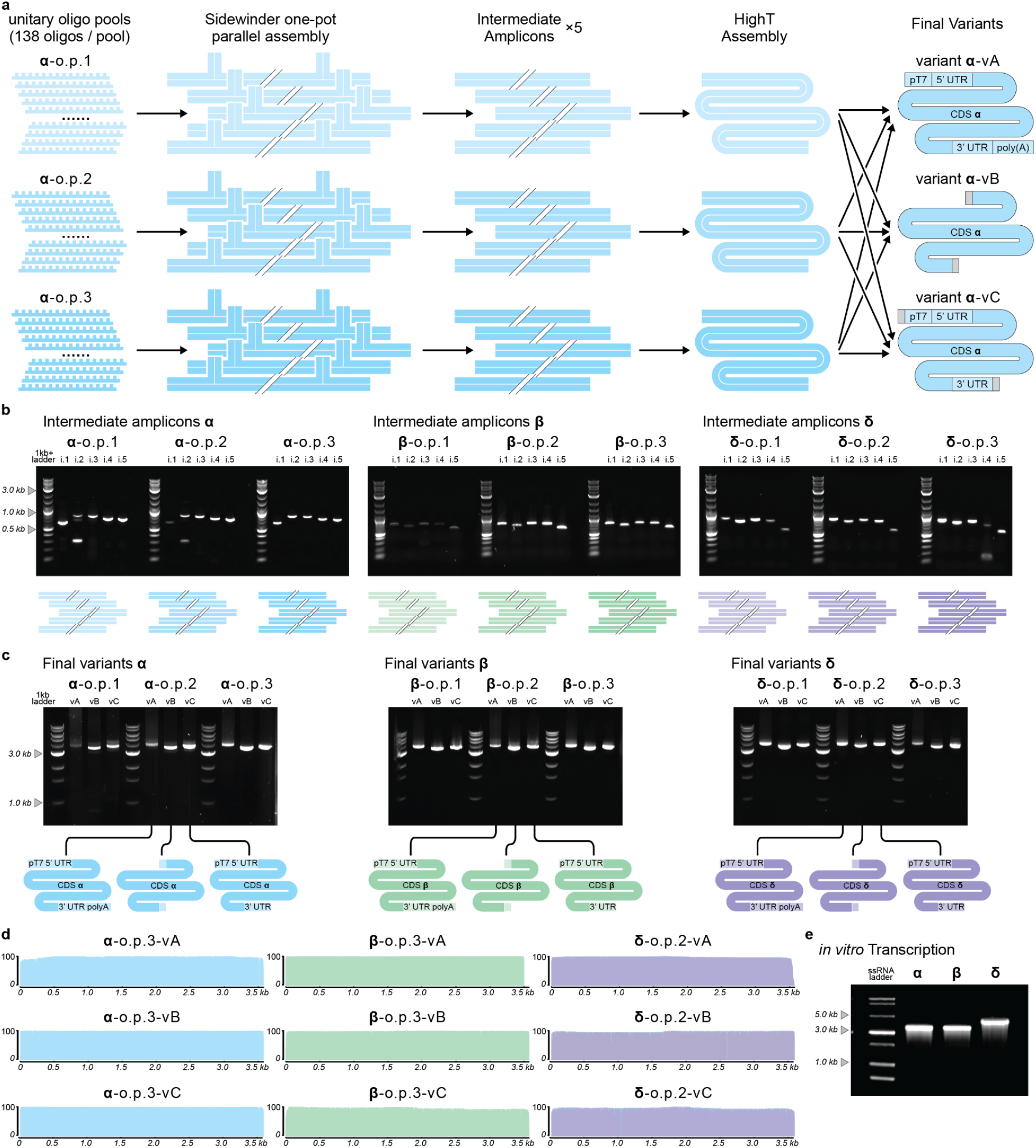
Sidewinder enables rapid and robust construction of long DNA products for mRNA production. **a)** Schematic overview depicting the molecular processing of DNA for CDS design α as a representative for all three CDS designs. Starting from unitary oligo pools (o.p.) with differing Sidewinder barcode designs, Sidewinder assembly is conducted producing the 3-way junction assembly for each pool in parallel. Intermediate PCRs are generated from the assembled pool. Entirely *in vitro*, the intermediate PCRs are then assembled into the full length product. Using this assembly as the template for a final PCR, different variant downstream products are generated which are used for *in vitro* transcription (vA), industry collaborator cloning (vB), or sequence verification cloning (vC). **b)** DNA agarose gel depicting 50 ng of the PCR products for each of the 5 intermediate amplicons for each of the 9 o.p. depicting a strong target band of the correct size in all instances. **c)** DNA agarose gel depicting 50 ng of the final PCR products for each of the 3 final variants (vA, vB, vC) for each CDS design (α, β, and δ) for each of the 9 o.p. depicting a single strong target band of the correct size in all instances. **d)** Coverage plots depicting Nanopore sequencing analysis with *de novo* assembly indicating high and consistent coverage for each DNA variant for one pool of each CDS design α (blue), β (green), and δ (purple). **e)** RNA gel depicting the gel extracted product of *in vitro* transcription for variant vA for each CDS design α, β, and δ.

Owing to the reliability of Sidewinder construction from oligo pools, the five intermediates from each of the nine o.p. were processed for assembly into the 3.75 kb target entirely *in vitro* without a cellular intermediate. Alternative methods of constructing large DNA products usually require an *in vivo* intermediate step where shorter DNA assemblies are conducted and subcloned into bacteria^22-25^ to physically isolate clones with correct constructs from a population with misassembled products. The *in vivo* intermediates ultimately increase cost and time owing to limited throughput and automation difficulty. Additionally, cell-free DNA construction is preferable in pharmaceutical applications due to concerns regarding byproducts such as endotoxins that persist after an *in vivo* subcloning step^26^.

After *in vitro* processing of the 45 intermediate amplicons, they were assembled in nine individual 5-piece assemblies. Using these final assembly products as the template for a PCR reaction enables us to process the final construct into three variants of the same core CDS design (**Fig. 2a**). The PCR for each variant (vA, vB, or vC) for each CDS design (α, β, δ) from each o.p. (o.p. 1, o.p. 2, o.p. 3) produced a prominent clean target band of the correct size in all cases (**Fig. 2c**). The amplicons for the first 9 successful variants (α_o.p. 3_ vA-vC, β_o.p. 3_ vA-vC, and δ_o.p. 2_ vA-vC) were imaged the morning of May 20th, resulting in 2 days between the delivery of oligos and the first successful construct for each CDS design (**Fig. 1a)**. The remaining 18 variants were all subsequently successfully constructed and imaged later that same day and into the next day, May 21st.

The first 9 successful variants were used for further validation and downstream processing. We used the samples for in-house Nanopore sequencing which had the run complete by the evening of May 20th, demonstrating a rapid and robust workflow for the construction and validation of DNA templates for mRNA vaccine candidates. Analysis of the sequencing results show *de novo* assembly aligning to the 3.75 kb target in each case with consistent and high coverage (**Fig. 2d**).

For each CDS design, variant vA is intended for use in downstream mRNA IVT and consequently the DNA template had a 120-base poly(A) tail added via PCR during the final PCR amplification step. Sidewinder’s high fidelity assembly enables incorporation of poly(A) tails in large DNA products constructed entirely *in vitro*. Although all constructs were verified to be correctly assembled, Nanopore’s inability to resolve long homopolymeric stretches^27^ can only qualitatively show the presence or absence of 3’ adenylation in the vA variants. To further assess the number of terminal adenine bases in the DNA template, we performed a restriction digestion on β_o.p. 3_ vA. Digestion with a specific cutter upstream of the poly(A) should produce a single 165-base fragment that includes the 120-base poly(A) tail. Running the digestion product on a TBE-PAGE gel shows a prominent band between 150-200 bases, indicating addition of the long poly(A) tail (**Extended data 1a**). Additionally, we conducted PCR to amplify the fifth intermediate amplicon both with and without the poly(A) in the primer. An agarose gel shows the expected migration distance consistent with the addition of the 120-base poly(A) tail (**Extended data 1b**). These data indicate reliable construction of the 3.75 kb vaccine candidate DNA template with reliable incorporation of a 120-base poly(A) tail entirely *de novo* and *in vitro*.

The variant vB for all candidates was designed to specifically amplify just the CDS component for downstream cloning and sequence verification by our industrial collaborator Centivax. The variant vC for all candidates was designed to specifically amplify the same template as vA but without the poly(A) tail to be used for in-house cloning into a plasmid.

Finally, α_o.p.3_ vA, β_o.p.3_ vA, and δ_o.p.2_ vA were used for IVT to validate the feasibility, robustness, and speed of in-house mRNA production via DNA templates from Sidewinder. RNA products were produced for each CDS variant with α and β yielding products of equivalent migration distance on an RNA gel and δ yielding an RNA product of slightly increased size (**Fig. 2e)**. This too was completed on May 20th, ultimately resulting in a completed workflow for construction, validation, and mRNA expression in just 2 days.

## Discussion

Viruses are a serious threat to public health. Many outbreaks are associated with zoonotic pathogen spillover to infect humans from an animal reservoir, and it has been predicted that this will become even more prevalent in the near future due to climate change^28^. A non-exhaustive list of viruses currently being monitored for epidemic potential includes coronaviruses, Ebola, and hantaviruses such as ANDV^29,30^. As the global response to COVID-19 recently demonstrated, our ability to quickly develop vaccines has improved in recent years, although even more rapid response will improve public safety measures and may more effectively reduce spread^31,32^. We have identified the production of template DNA as a key bottleneck in the development process of mRNA vaccines. Although DNA construction is just one aspect of the complex process of mRNA vaccine development, relieving this bottleneck via Sidewinder enables focus on downstream vaccine study and development to improve safety and efficacy. Towards this end, the DNA designs and constructs produced in this study are available for researchers worldwide to use for mRNA vaccine validation against the ANDV infection.

Applying Sidewinder and our novel design algorithm PyWinder, we were able to accelerate the development process for mRNA vaccine candidates via rapid production of DNA templates. The entire process (identification of a viral antigen, design of DNA, delivery of oligo pools, assembly of DNA template, sequence verification of the constructed DNA template, and in vitro transcription to manufacture mature mRNA with cap and polyadenylation) required only 2 weeks, with just 2 days of in silico design and 2 days of in physico DNA construction, validation, and mRNA production upon oligo pool delivery. Using Sidewinder for assembly of DNA templates helps alleviate a major bottleneck in rapid mRNA vaccine development and is performed entirely *in vitro*, bypassing conventional time-consuming *in vivo* cloning workflows entirely.

The speed of Sidewinder enabled our multifaceted approach to design and construction. By implementing multiple coding sequence designs in α, β, and δ, we may better protect against delays in development caused by particular codon choices influencing large scale production or immunogenicity. We observed differences in migration pattern on the nondenaturing - RNA gel following IVT for α, β, and δ CDS designs, potentially indicating mRNA secondary structure differences between codon choice. Further, by designing multiple oligo pools for each CDS design, we better protect against delays in development caused by any failure points that may have arisen in oligo synthesis, DNA construction, or DNA amplification. Ultimately, all 9 oligo pool designs successfully produced the intended 3.75 kb target and all were further processed into each of the three variant applications vA, vB, and vC. By using a single construct template for multiple PCR reactions, we generated vA for α, β, and δ which was transcribed with IVT into mRNA. We generated vB for α, β, and δ which was designed to be sent to Centivax for cloning and downstream development. We generated vC for α, β, and δ which was designed for cloning in-house for reliable storage and replication of sequence-verified DNA templates for preservation and distribution of the mRNA vaccine candidate templates.

In the present work, we focus entirely on the construction and transcription of upstream candidate constructs that may be further interrogated for delivery, safety, and efficacy. We anticipate that the described workflow enabled by Sidewinder DNA assembly can be applied to accelerate the development and testing of mRNA vaccines against a variety of pathogens and cancers. This could be pertinent for accelerating the manufacture of personalized cancer neoantigen vaccines, an application which is particularly sensitive to construction delays^33^.

As the design of DNA- and RNA-based medicines advances at a rapid pace, we show in this work that Sidewinder is a tool to realize and accelerate the development of these next-generation therapies, with the goal of improving patient outcomes for a myriad of diseases.

## Methods

### Choosing the Protein Sequence

To analyze the likelihood of M polyprotein antigen MHC display, we employed NetMHCpan-4.1 and NetMHCIIpan-4.1 with peptide fragments generated by the Immune Epitotope Database & Tools (IEDB) website (https://nextgen-tools.iedb.org)^1,2^. For the peptide-MHC Class I presentation analysis (NetMHCIpan-4.1 EL), we constrained prediction to a peptide size of 8-11 residues and 27 common HLA types (covering ~97% of the human population). Out of the 121,986 possible peptides that were analyzed, 910 were predicted to be strong MHC Class I binders (NetMHCIpan_EL %Rank < 0.5%). For the peptide-MHC Class II presentation analysis (NetMHCIIpan-4.1 EL), we constrained prediction to a peptide size of 15-25 residues, the 7 most common HLA types (covering ~50% of the human population), and a peptide shift length of 5. Out of the 17,262 possible peptides that were analyzed, 101 were predicted to be strong MHC Class II binders (NetMHCIIpan_EL %Rank under 2%). This indicates that M polyprotein antigens have a high predicted likelihood of being presented to CD8+ Cytotoxic T-cells and CD4+ Helper T-cells, thereby triggering an immune response in a human recipient.

### mRNA sequence design

The overall design of our mRNA vaccine constructs consisted of five major components: a T7 promoter, a 5′ untranslated region (5′ UTR), an antigen-coding sequence (CDS), a 3′ untranslated region (3′ UTR), and a poly(A) tail. The T7 promoter sequence was designed to be compatible with the New England Biolabs *in vitro* transcription and 5’ capping system (New England Biolabs: HiScribe® T7 mRNA Kit with CleanCap® Reagent AG, E2080S) to ensure efficient RNA synthesis.

The 5′ UTR and 3′ UTR sequences for each vaccine construct were computationally designed using the GEMORNA platform^3^. Specifically, the 5′ UTR generation and 3′ UTR generation modules were used to generate candidate untranslated regions of varying lengths, including five short, ten medium-length, and five long candidates for each region. Following sequence generation, all candidate UTRs were re-evaluated using the corresponding 5′ UTR and 3′ UTR prediction modules within GEMORNA to estimate their predicted translational performance and stability. The highest-scoring UTR sequences were subsequently selected as the design for downstream vaccine construct assembly.

Based on the amino acid sequence of the ANDV M polyprotein antigen (Genbank Accession: QDI78308.1), three distinct CDS variants were designed, designated α (alpha), β (beta), and δ (delta). The α variant was generated using the CDS optimization module of GEMORNA to improve predicted translational performance. The β variant consisted of a separately optimized coding sequence that was generously provided by our industrial collaborator, Centivax. In contrast, the δ variant retained the native viral coding sequence obtained from GenBank (Genbank Accession: MN095801.1), serving as a non-optimized reference construct.

A poly(A) tail of 120 nucleotides was added via PCR in variant vA of each construct, consistent with commonly used poly(A) tail lengths in mRNA vaccine design to support transcript stability.

### Oligo design

Sidewinder oligos were designed using either the original or modified versions of PyWinder^4^ with thermodynamic filtering. In the modified versions, only the barcode generation and thermodynamic filtering steps of the algorithm were edited.

For the α_2_, α_3_, β_2_, β_3_, and δ_2_, δ_3_ strands, the Sequence-Symmetry Minimization^5,6^ step’s default pick of *q* and *k* parameters discussed previously^4^ (the maximum length of a shared substring between the picked toehold set and the barcode set, and the maximum length of a shared substring within the barcode set respectively) was modified such that *q* ← *q* + 1 and *k* ← *k* + 1. For δ_1_, due to sequence complexity, we also modified default *q* and *k* such that *q* ← *q* + 2 and *k* ← *k* + 2. The α_1_ and β_1_ strand designs used the default *q* and *k* parameters.

PyWinder also includes a thermodynamic-based filtering system, which we used to make all oligo pool designs (o.p.d.). PyWinder’s primarily string-based approach is advantageous over other predominantly thermodynamic approaches for orthogonal barcode design because explicit thermodynamic design criteria are computationally expensive (and can overfocus on thermodynamic optimization without kinetics consideration)^7^. However, because calculated physical thermodynamic properties may provide valuable information beyond string similarity, we nonetheless aimed to incorporate such properties into the design process, albeit cheaply. Specifically, we applied a thermodynamic-based filtering mechanism across a smaller set of string-generated designs to decrease thermodynamic computation cost.

Under our thermodynamic filtering, the full string pipeline is first run *M* times to create *M* Sidewinder strand sets. Each of these sets is saved and analyzed thermodynamically. This analysis is as follows: for each string-generated library, we extract the concatenated toehold-barcode strands *z*_*i*_ = *τ*_*i*_ • β_*i*_ and compute a full duplex pairwise off-target probability matrix *P*^*off*^. To calculate *P*^*off*^, let *E*([(*a, b*)] | *I*_*c*_, *T*, Θ, Ψ) denote the expected concentration of (*a, b*) (which represents all DNA complexes composed solely of a single strand of *a* and a single strand of *b*) where *I*_*c*_ is the set of any non-zero initial concentrations for starting species, *T* is the temperature, Θ is the the set of salt conditions, and Ψ is the partition set of complexes considered possible to form from the initial species. Any complex not in Ψ will not be considered in intermediate calculations and so will have no effect on returned final concentrations. Consequently, the choice of complexes in Ψ represents an assumption of which complexes are thermodynamically accessible and significant (i.e. those ∈ Ψ) and which are insignificant (i.e. those ∉ Ψ).

Then, letting 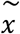 denote the reverse complement of DNA sequence *x*, and defining Γ_*xy*_ = (*I*_*c*_ = *i*_*xyc*_, *T* = *t*, Θ = θ, Ψ = ψ_*xy*_) and *c* = 10^−8^ *M, i*_*xyc*_ = {[*x*] = *c*, [*y*] = *c*}, *t* = 50° *C*, θ = {[*Na*^+^] = 0. 154*M*, [*Mg* ^2+^] = 0. 01*M*}, and ψ_*xy*_ = {(*x*), (*y*), (*x, x*), (*y, y*), (*x, y*)}, we write 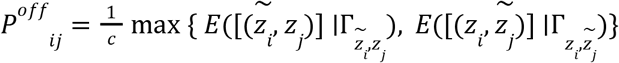. Our choice of 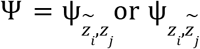 in calculating 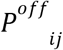 means we are only considering coding-sense strand to template-sense strand toehold-barcode *off*-target interactions (since these are the interactions in Sidewinder assembly at a higher risk of mis-ligation). It also means that we neglect consideration of any complexes involving more strands than a duplex, reducing computational cost. Any *E*([(*a, b*)] | *I*_*c*_, *T*, Θ, Ψ) calculations were completed using the NUPACK 4.0.2.0 thermodynamic algorithm^8,9^.

Next, each of the *M* string-generated libraries is given a score 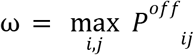. All *M* string-generated libraries are then ranked (with a lower ω implying a better library design). Then, CDS design (α, β, δ) o.p.d. 1 corresponds to the top ranked library in an *M* = 10 regime, and o.p.d. 2 and o.p.d. 3 correspond to the 1st and 2nd ranked libraries in an *M* = 1000 regime respectively.

### Oligo Purchasing

Oligo pools were purchased from Integrated DNA Technologies (IDT) as oPools™ Oligo Pools at the 50 pmol/oligo synthesis scale. PCR amplification primers were also ordered from IDT and were purified by standard desalting treatment and shipped dry. All oligos are listed in **Supplementary Table 1**.

### Oligo phosphorylation and annealing

Oligo pools were resuspended in 100 µL of 1× T4 DNA Ligase Reaction Buffer (New England Biolabs: B0202S) and gently vortexed. A 5.4 µL aliquot of the oligo pool was taken for phosphorylation in a 10 µL reaction of 1× HiFi Taq DNA Ligase Reaction Buffer (New England Biolabs: B0647SVIAL) with 1.1 µL of T4PNK (New England Biolabs: M0201S) at 37°C for 1 hour on a thermocycler. Immediately after phosphorylation, the thermocycler temperature was raised to 98°C for enzyme deactivation and oligo denaturation. The temperature was then decreased at −1°C per minute to 25°C and held at 25°C until use.

### Sidewinder Assembly

Assembly reactions were conducted in 50 µL final volume of 1× HiFi Taq DNA Ligase Reaction Buffer. The entire 10 µL phosphorylated, annealed oligo pool was added to 5 µL of 10× HiFi Taq DNA Ligase Reaction Buffer (New England Biolabs: B0647SVIAL) and 33 µL of Milli-Q® water. The 48 µL mixture was placed on the thermocycler and 2 µL of Taq ligase (New England Biolabs: M0647S) was added after the reaction reached the intended temperature on the thermocycler. Thermocycler protocol was as follows:

1. 68°C for 5 minutes.
2. Add ligase.
3. Drop temperature from 68°C to 37°C at −1°C per minute.
4. Hold at 37°C for 20 minutes.
5. Raise temperature back to 68°C.
6. Repeat steps 3-5 for a total of 10 cycles.
7. Hold at 37°C for 8 hours.

### PCR amplification and purification of intermediate products

As described, CDS designs α, β, and δ each had 3 Sidewinder oligo pool designs. For each of the 9 ligated Sidewinder assemblies, repliQa HiFi ToughMix (Quantabio: 95200-025) was used to amplify segments of the 3WJ assembly with deoxyuridine (dU) containing primers. These intermediate amplicons were approximately 0.7-1 kb which together span the length of the ~4 kb target construct. We observed that 1 μL of unpurified three-way junction (3WJ) assembly from the previous assembly step was sufficient template in a 50 μL PCR reaction for robust amplification. PCR reaction conditions were established according to manufacturer recommendations and predicted Tm of primers on SnapGene.

Thermocycler settings for PCR:

α intermediate PCRs

1. 98°C for 45 seconds.
2. 98°C for 15 seconds.
3. 68°C for 6 seconds.
  a. Go to Step 2 for 40 cycles.
4. 68°C for 5 minutes.

β and δ intermediate PCRs

1. 98°C for 45 seconds.
2. 98°C for 15 seconds.
3. 63°C for 5 seconds.
4. 68°C for 5 seconds.
  a. Go to Step 2 for 40 cycles.
5. 68°C for 5 minutes.

α, β, δ final assembly PCRs

1. 98°C for 45 seconds.
2. 98°C for 15 seconds.
3. 63°C for 5 seconds.
4. 68°C for 20 seconds.
  a. Go to Step 2 for 40 cycles.
5. 68°C for 5 minutes.

After each product was amplified, the 50 µL of PCR product was purified using a QIAquick PCR Purification Kit (Qiagen: 28104) and eluted in 40 µL of 10% elution buffer. Amplicons with low yield of the correct size product had multiple 50 µL reactions passed through the same column. Concentration of purified product was measured using the Qubit 1x dsDNA High Sensitivity Assay Kit (Invitrogen, Thermo Fisher Scientific: Q33230).

### Hierarchical construction

For all conditions, the purified (Qiagen: 28104) intermediate amplicons were assembled *in vitro* with HighT USER assembly into the final size of ~4 kb. 30 µL of the purified intermediate amplicons were first digested at 37°C in a 40 µL reaction in 1× rCutSmart® with 1.1 µL of USER® enzyme cocktail (New England Biolabs: M5505S) to expose assembly overhangs via removal of the deoxyuridine (dU). The digests were purified via gel extraction (New England Biolabs: T1120S) and assembled at 8 nM in a 50 µL reaction of 1× HiFi Taq DNA Ligase Reaction Buffer at 50°C for 16 hr with 2 µL Taq ligase. After assembly, 1 µL of the reaction mixture was used as the template in a 50 µL PCR. This template was used for each of the downstream amplifications to produce the final variants vA, vB, and vC.

### Variant vA: *in vitro* transcription

Variant vA products were designed to serve directly as linear DNA templates for *in vitro* transcription (IVT). Following Sidewinder DNA assembly, the final constructs corresponding to the CDS designs α, β, and δ were amplified via PCR using a reverse primer containing a 120-base poly(T) sequence to incorporate a poly(A) tail into the resulting DNA templates.

The amplified products were subsequently resolved by agarose gel electrophoresis, and bands corresponding to the expected sizes were excised and purified by gel extraction (New England Biolabs: T1120S). To further remove residual proteins, enzymes, and other potential contaminants that could interfere with downstream IVT reactions, purified DNA products were subjected to phenol–chloroform extraction followed by ethanol precipitation (**Methods**).

Purified linear dsDNA templates were then used for *in vitro* transcription using the HiScribe T7 mRNA Kit with CleanCap AG (New England Biolabs: E2080S). IVT reactions were performed according to the manufacturer’s recommended conditions. The resulting RNA products were analyzed by gel electrophoresis using the EMBER Ultra RNA Gel Kit (Biotium: #41044-T) to assess transcript purity and length. After the RNA products were visualized depicting prominent bands of the expected size, we used the Monarch® Spin RNA Cleanup Kit (New England Biolabs: T2040S) to extract the corresponding band, demonstrating isolation and purification of intended mRNA products depicted in the gel with ssRNA ladder (New England Biolabs: N0362S) on **Fig 2e**.

### Variant vB: Centivax CDS design for cloning/manufacturing

After construction, just the CDS region of each full assembly (α_1-3_, β_1-3_, and δ_1-3_) was amplified with vB primers that added restriction sites for downstream use in Centivax’s internal mRNA production platform.

### Variant vC: Sequence verified template for global distribution

Variant vC was designed for incorporation into a plasmid and contained a T7 promoter, 5’UTR, CDS and 3’UTR without the poly(A) tail. The ultimate intent for this sequence is for the potential distribution of a sequence-perfect plasmid to researchers worldwide if the need arises.

### Phenol-chloroform extraction

To perform phenol-chloroform extraction, phase separation of DNA is initiated by adding an equal volume of 1:1 phenol/chloroform mixture, or 25:24:1 phenol:chloroform:isoamyl alcohol (Sigma-Aldrich: P2069-400ML) and vortexed 5-10 seconds or inverted thoroughly to mix. After centrifuging at 12,000-16,000 × g for 5 min at room temperature, we carefully transferred the upper aqueous phase (~90% of volume) to a new RNase-free tube. We then extracted twice with an equal volume of chloroform to remove residual phenol. Afterwards, we precipitated the DNA by adding 1/10 volume of 3 M sodium acetate, pH 5.2 (Macron Fine Chemicals: 7364-04), and two volumes of ethanol (Koptec: 64-17-5, diluted as needed with Milli-Q® water) and incubating it at −20°C for 2-4 hours. We then pelleted the DNA in a microcentrifuge for 15 minutes at 16,000 × g at 4°C and removed the supernatant. After rinsing the pellet by adding 500 μL of 70% ethanol and centrifuging for 15 minutes at 16,000 × g, we discarded the supernatant. Finally, we air dried the pellet for 5-10 minutes and resuspended it in nuclease-free water at room temperature for 20 minutes.

### DNA gel imaging

Gel electrophoresis was conducted with 1-2% agarose gels (Affymetrix: 9012-36-6) stained with SYBR™ Safe (Invitrogen, Thermo Fisher Scientific: S33102) in 0.5× TBE buffer (Genesee Scientific: 20-196) run for 25-40 minutes at 135 volts. The DNA ladder used were Low Molecular Weight DNA Ladder (New England Biolabs: N3233S), 1 kb DNA Ladder (New England Biolabs: N3232S), and 1 kb Plus DNA Ladder (New England Biolabs: N3200S) and labeled where used. Gels were imaged using BioRad ChemiDoc Imaging System (BioRad: 12003153). Main figure gels show 50 ng of template loaded, as measured using the Qubit 1x dsDNA High Sensitivity Assay Kit (Invitrogen, Thermo Fisher Scientific: Q33230).

The TBE-PAGE gel depicted in **Extended data 1a** was conducted with Novex TBE 8% 15-well gel (Invitrogen, Thermo Fisher Scientific: EC62155BOX) run at 200 volts for 30 minutes and stained with 5 µL of SYBR™ Safe in 25 mL water for 10 minutes with gentle rocking.

### Sequencing library preparation

DNA libraries were prepared using the Oxford Nanopore Technologies (ONT) Native Barcoding Kit 24 V14 (SQK-NBD114.24) following the manufacturer’s instructions. Briefly, PCR amplification products underwent end-repair and dA-tailing, followed by native barcode ligation and pooling of barcoded samples. Sequencing adapters were subsequently ligated prior to library cleanup and loading onto a MinION Mk1B device equipped with an R10.4.1 flow cell (FLO-MIN114). Sequencing was performed using MinKNOW with super high-accuracy basecalling enabled.

### Sequencing analysis

Sequencing reads were basecalled and demultiplexed, and adapter/barcode sequences were removed. Filtered ONT reads were assembled de novo with Flye v2.9.6 using the ONT high fidelity read mode, followed by consensus polishing with Medaka v2.2.1 using the model appropriate for the flow cell and basecaller configuration. Polished assemblies were used as sample-specific references. Reads were mapped back to the polished assembly with minimap2^10,11^, alignments were sorted and indexed with SAMtools^12^, and per-base pileups and depth of coverage were generated using samtools mpileup/samtools depth to assess read support and coverage across the assembled reference.

## Acknowledgements

We thank Dr. J. Merson for helpful discussion on the work.

We thank Dr. A. Woolfson and Mr. J. Smith for their helpful discussion when formulating the project.

## Author contributions

K.W. conceived the study. J.A., A.D, S.K., and J.G. identified the protein target and designed the CDS design β. J.S.P. designed the Sidewinder oligos; N.E.R., W.Z., S.W., T.Z., E.A. and D.G. performed DNA assembly experiments. T.Z. performed IVT on the products. T.G., J.H., J.S.H., C.S., R.Z., and J.Z., performed sequencing and analyzed results. All of the authors discussed the results. The manuscript was written with input from all authors.

## Additional information

Supplementary information is available for this paper. Correspondence and requests for materials should be addressed to.

## Data availability

The DNA products synthesized in this paper will be made available upon request. The oligos used in this study are listed in **Supplementary Table 1** and the CDS design variants are listed in **Supplementary Table 2**.

## Extended Data figures

**Extended Data 1.**
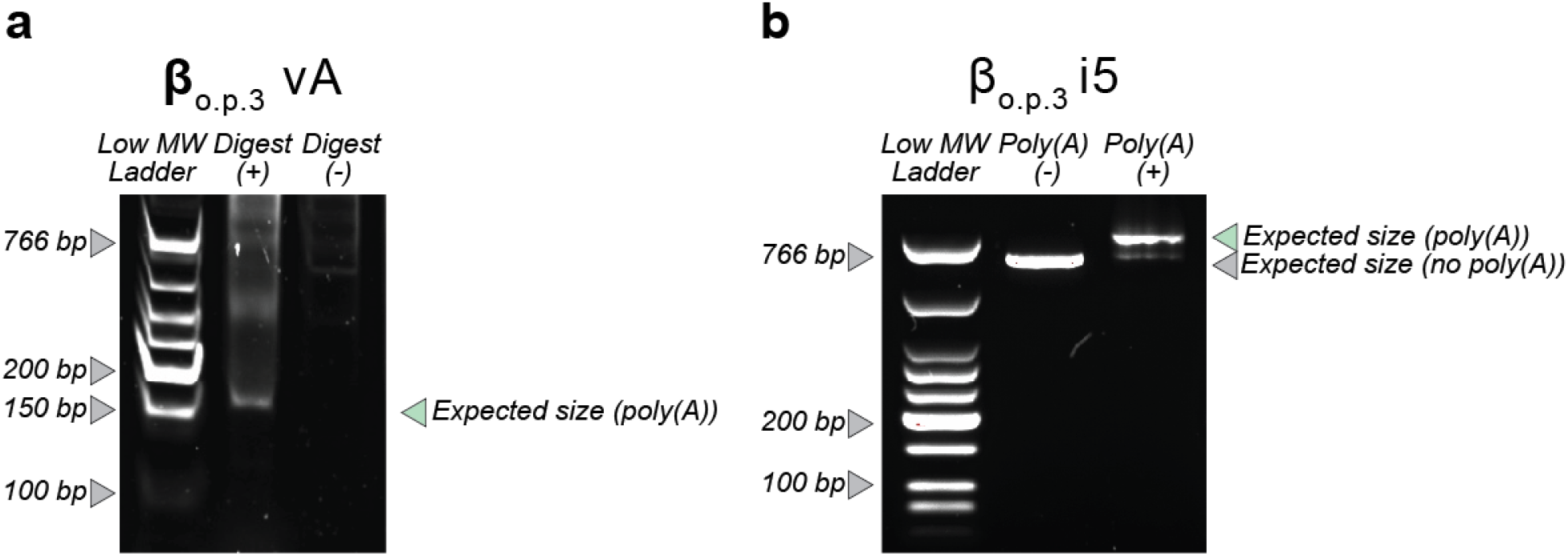
Validation of poly(A) tail incorporation and size. **a)** TBE-PAGE gel depicting enzymatically digested (+) and undigested (-) β-o.p.3-vA near the poly(A) site. The digested product should be 165 bases and shows a band at the expected size, suggesting the presence of the correctly sized poly(A) tail. **b)** DNA agarose gel depicting 100 ng of the β-o.p.3 5th intermediate PCR product with and without the 120-base poly(A) tail. Both reactions show a prominent band of the correct size, 655 bp without the poly(A) tail (-) and 775 bp with the poly(A) tail (+), consistent with the incorporation of the 120-base poly(A) tail.

